# Evidence of positive selection and a novel phylogeny among five subspecies of song sparrow (*Melospiza melodia*) in Alaska

**DOI:** 10.1101/2024.05.21.595201

**Authors:** Caitlyn C. Oliver Brown, Keiler A. Collier, Kendall K. Mills, Fern Spaulding, Travis C. Glenn, Christin L. Pruett, Kevin Winker

## Abstract

Under local adaptation, populations evolve traits in response to the local environment. Isolated island populations often experience different selection pressures than their mainland counterparts, which enables the study of how phenotypes and genotypes respond to different selection regimes. We used a group of five phenotypically differentiated subspecies of song sparrow (*Melospiza melodia*) in Alaska to examine the effects of local adaptation. Song sparrows occur across southern Alaska from Attu Island in the western Aleutian Islands, to southeast Alaska. Moving from western to eastern Alaska these populations demonstrate striking body size differences (larger-to-smaller) and a change from a sedentary to a migratory/partially migratory life-history strategy. We examined the phenotypic attributes of these populations and used whole-genomic data to determine relationships and test candidate loci for evidence of selection. Phenotypic measurements of museum specimens (n = 227) quantified the dramatic size differences among these populations, with westernmost *M. m. maxima* being ∼1.6 times larger than easternmost *M. m. rufina*. Ultraconserved elements (UCEs) were extracted for phylogenetic reconstruction and candidate genes were extracted for selection testing. We analyzed 26 candidate genes for body size, migration and dispersal, color, and salt tolerance. Two of the candidate genes showed signs of positive selection: *BCO1* (associated with plumage color) and *KCTD21* (associated with dispersal). Phylogenetic analysis of UCEs showed *M. m. maxima* as sister to the other Alaska *M. melodia* subspecies. This suggests *M. m. maxima* colonized earliest, perhaps before the last glacial maximum, and that Alaska was later recolonized by ancestors of the remaining four subspecies.

## Introduction

Our current understanding of evolution suggests that adaptation to local conditions enables populations to persist and that geographic variation among populations often reflects these processes. Local adaptation is defined as the process by which a local population evolves traits best suited for its environment (Turesson, 1922; Mayr, 1963; Williams, 1966; Kawecki & Ebert, 2004; Savolainen, Lascoux & Merilä, 2013). Local adaptation to a specific environment can potentially lead to speciation (Savolainen, Lascoux & Merilä, 2013; Tiffin & Ross-Ibarra, 2014), but this process of adaptation can be hindered by the homogenizing effects of dispersal and gene flow (Kawecki & Ebert, 2004; Savolainen, Lascoux & Merilä, 2013). While it is still challenging to identify the genetic basis of local adaptation, the advent of high-throughput genomic data can be used to identify potential genes under selection (Tiffin & Ross-Ibarra, 2014; Bomblies & Peichel, 2022). Phenotypically differentiated island populations can be used to examine the genetic basis of local adaptation. This has been done in many terrestrial organisms, such as lizards (Losos, Warheitt & Schoener, 1997), birds (Grant, 1981), spiders (Gillespie, 2002), and plants (Choi et al., 2021).

Island systems have been influential in our understanding of the processes of evolution (MacArthur & Wilson, 1967; Losos & Ricklefs, 2009; Warren et al., 2015). Islands, especially archipelagos, are natural laboratories to study evolution, given their relatively young geological age and geographic isolation from the mainland (Losos & Ricklefs, 2009; Warren et al., 2015). Because of these conditions, island populations provide a way to determine how local adaptation accrues. The majority of research on local adaptation has occurred on tropical or mid-latitude islands, however there has been a recent focus on high-latitude island systems (Whittaker & Fernadez-Palacios, 2007). For example, the Aleutian Islands in the North Pacific Ocean constitute a volcanic archipelago that extends ∼1,800 km from western Alaska towards eastern Russia (Murie, 1959). During the Pleistocene, Alaska experienced multiple cycles of glaciation and glaciers covered much of the continent. However, ice-free zones, or refugia, were present which could have allowed populations to persist and the Aleutian Islands are hypothesized to contain some of these refugia (Pruett & Winker, 2005a; Winker et al., 2023). Isolated avian populations on these islands often exhibit both lower genetic diversity than mainland counterparts and phenotypic differences, such as larger body size (Pruett, Li & Winker, 2018). One such species is the song sparrow (*Melospiza melodia*, Passeriformes: Passerellidae), a widely distributed songbird found only in North America with ∼25 recognized subspecies (Aldrich, 1984; Patten & Pruett, 2009). These subspecies exhibit a wide range of phenotypes and life histories and reside in a variety of environments across North America (Arcese et al., 2020; Carbeck et al., 2023). Because of this, song sparrows are an excellent candidate for the study of local adaptation.

Five song sparrow subspecies occur across southern Alaska, from the western Aleutian Islands to southeast Alaska. These subspecies, from west to east, are *M. m. maxima, M. m. sanaka, M. m. insignis, M. m. caurina*, and *M. m. rufina* (Fig. 1; Gibson & Withrow, 2015). Prior genetic research suggested that ancestral song sparrows colonized the Aleutian Islands sequentially, from east to west (Pruett & Winker, 2005b). Across this complex, these subspecies exhibit different life history traits, presumably due to differences in their environments. The subspecies that occur furthest west (*maxima* and *sanaka*) are non-migratory residents on the Aleutian Islands, a relatively harsh maritime environment. These two subspecies are found in grassy slopes along marine beaches, a habitat considered marginal for continental subspecies (Murie, 1959; Aldrich, 1984). The subspecies that occur furthest east in Alaska (*caurina* and *rufina*) are seasonal migrants, residing largely on continental North America and migrate south for the winter (Aldrich, 1984; Patten & Pruett, 2009). One subspecies (*insignis*) is primarily a year-round resident of the Kodiak Archipelago (Patten & Pruett, 2009). There are obvious phenotypic and natural history differences among these subspecies. The most notable phenotypic characteristics are differences in plumage coloration and body size, wherein the most western subspecies in the Aleutian Islands are much larger than the eastern subspecies in southeast Alaska (Aldrich, 1984; Pruett & Winker, 2005b; Patten & Pruett, 2009). These phenotypic differences suggest these five subspecies have adapted to their local environments (Fig. 2; Winker, 2010).

**Figure 1:**
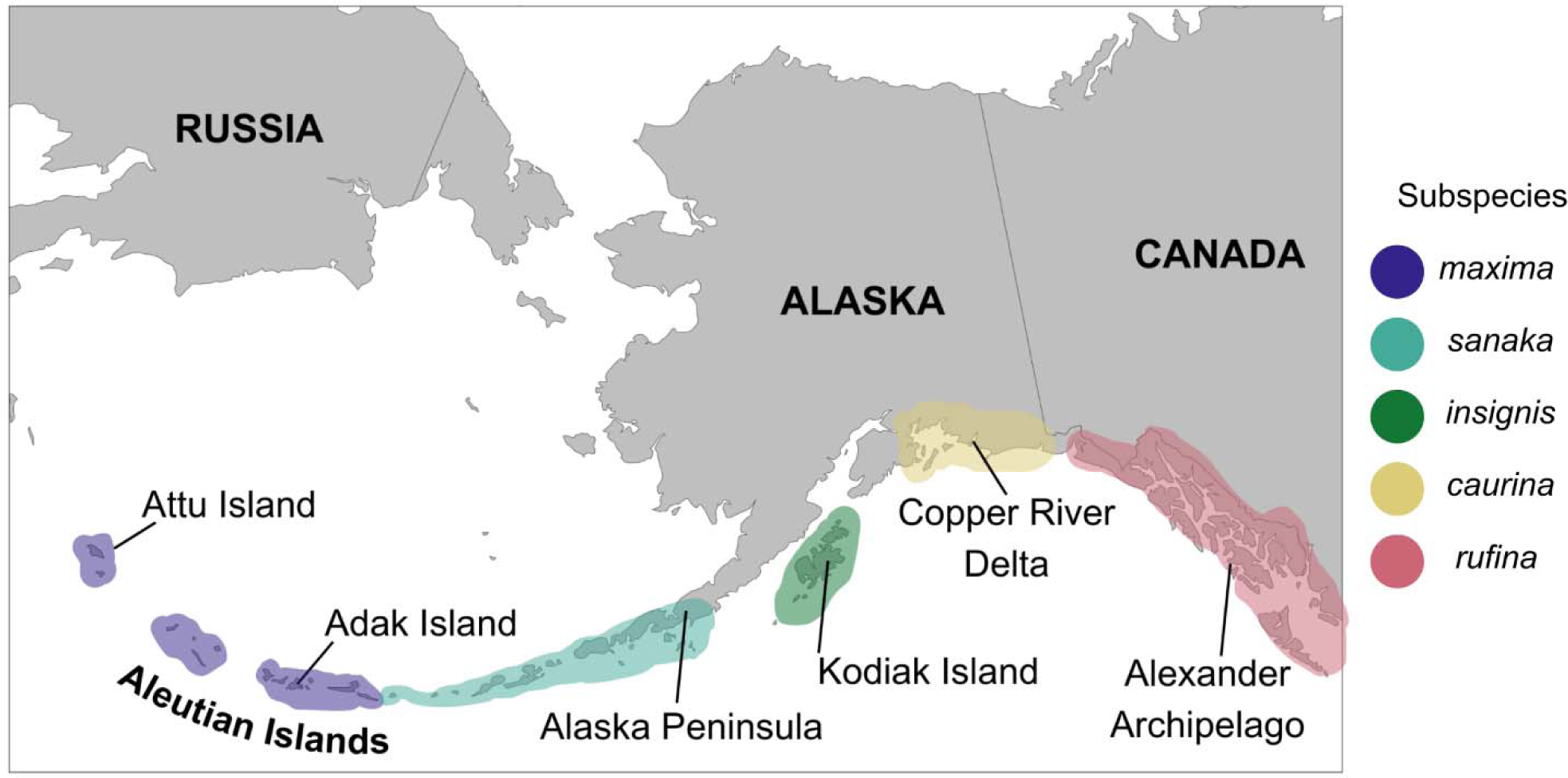
Distribution of five subspecies of song sparrow (*Melospiza melodia*) in Alaska. From west to east: *M. m. maxima* (dark blue), *M. m. sanaka* (teal), *M. m. insignis* (green), *M. m. caurina* (yellow), and *M. m. rufina* (pink).

**Figure 2:**
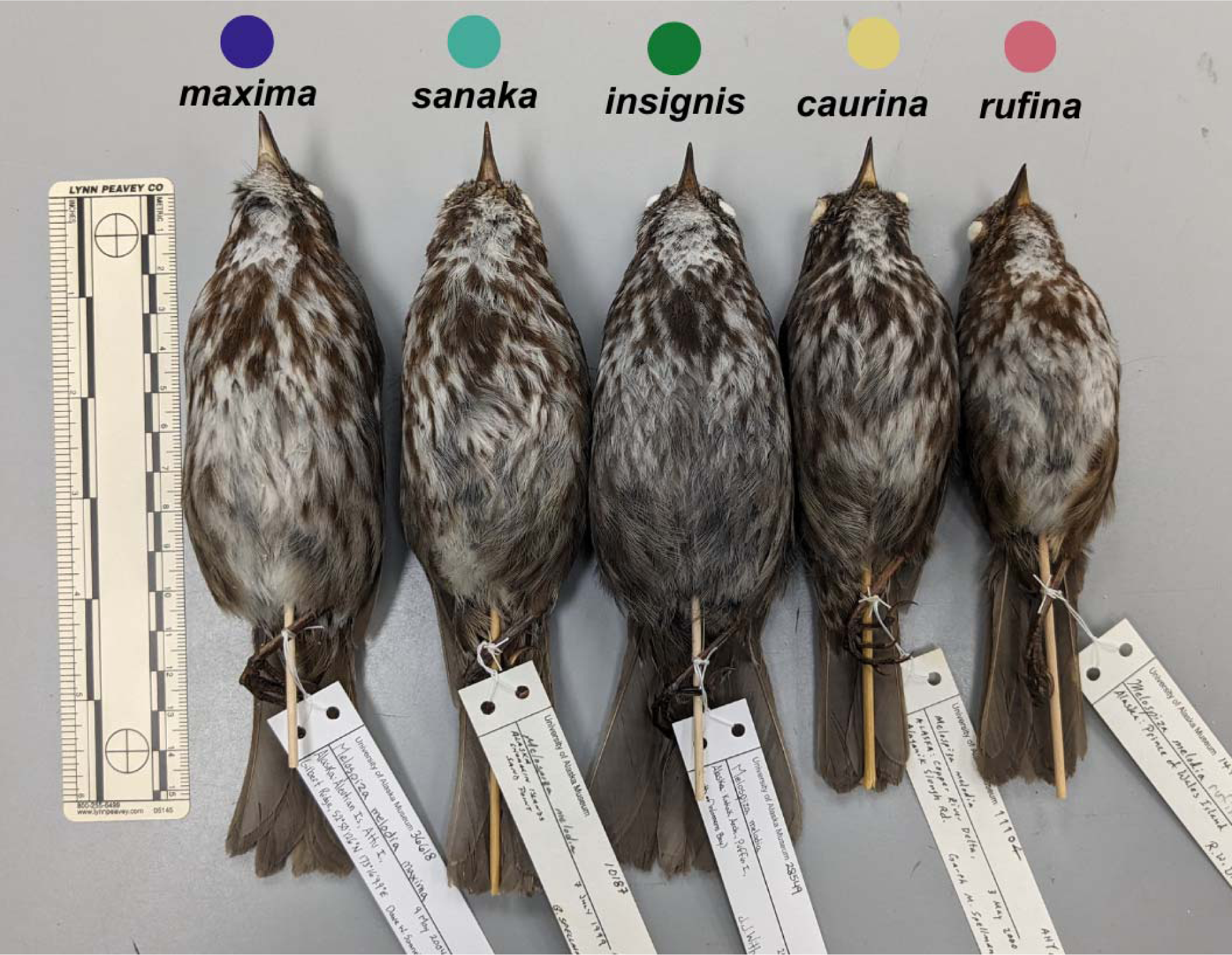
Museum specimens of five *M. melodia* subspecies of interest that occur in Alaska. Colored circles correspond to the colors in Figure 1. Photo credit: Caitlyn C. Oliver Brown

We studied phenotypic differentiation relevant to aspects of life history characteristics among the subspecies of the Alaska song sparrow complex to examine adaptive evolution of high-latitude island populations. We used a combination of genomic and morphological analyses using museum specimens. We asked three questions. First, which morphological traits are significantly different among the five *M. melodia* subspecies in Alaska? Second, are candidate genes, associated with key phenotypic attributes, under positive selection in the most extreme variant, *M. m. maxima*? Third, what are the phylogenetic relationships among these subspecies?

## Materials and Methods

### Phenotypic Analysis

To examine the phenotypic variation among the five *Melospiza melodia* subspecies, we obtained data from 227 vouchered specimens from the University of Alaska Museum (Table S1). We restricted our analyses to adult males with complete phenotypic and locality data. Phenotypic data included: mass, wing chord length, tail length, tarsus length, bill length, bill height, bill width, and skull length (Winker, 2000). Measurements were taken to the nearest 0.1 millimeter (mm) or to 0.1 gram (g) for mass prior to specimen preparation (Winker, 2000). Data were reviewed and analyzed using R (v. 4.1.1; R Core Team, 2021) and RStudio (v. 2022.12.0.353; Posit Team, 2022) and accompanying packages, “devtools” (v.2.4.5; Wickham et al., 2022) and “tidyverse” (v.1.3.2; Wickham et al., 2019). We removed outliers that were three standard deviations away from the mean to account for human measurement error. Phenotypic data were then visualized with boxplots and a principal components analysis (PCA) using the packages “ggplot2” (v.3.4.1; Wickham et al., 2019), “ggpubr” (v. 0.6.0; Kassambara, 2023), and “ggord” (v. 1.1.7; Beck, 2022). ANOVA tests were used for each measurement and pairwise Tukey post-hoc tests were used to determine which subspecies pairs were different. R scripts and raw phenotypic data are available at https://github.com/coliverbrown/melospiza-melodia-phenotype.

### Sampling and Laboratory

For genomic analysis, we sampled high-quality vouchered tissue samples from wild individuals archived at the University of Alaska Museum (Table S2). Our sample size included one individual from each of the five *M. melodia* subspecies that occur in Alaska. Two taxa were included as outgroups: the swamp sparrow (*M. georgiana*) and the Lincoln’s sparrow (*M. lincolnii*). DNA extractions followed standard protocol for animal tissues using the QIAGEN DNeasy Blood + Tissue Extraction Kit (Valencia, California).

Libraries were prepared using the iTru (Illumina dual-index) library protocols described in Glenn et al. (2019). Briefly, we sheared the genomic DNA using a Bioruptor (Diagenode, Denville, NJ, USA) targeting a 500bp average fragment size. The sheared DNA was end-repaired, adenylated, and ligated to iTru adapters followed by limited-cycle PCR of iTru primers to add indexes and complete the library molecules using Kapa Library Preparation Kit reagents (Kapa Biosystems [Roche, Basel, Switzerland]). We sequenced the pooled libraries on an Illumina sequencer (Illumina, San Diego, CA, USA) to obtain paired-end (PE) 100 base reads.

### Bioinformatics

After sequencing, we constructed whole genome assemblies for each of the sequenced individuals. Our bioinformatics pipeline centered on the package PHYLUCE (Faircloth, 2016). Raw and untrimmed FASTQ data that contained low-quality bases were removed using Illumiprocessor (Faircloth, 2013), which incorporates Trimmomatic (Bolger, Lohse & Usadel, 2014). Raw paired end reads (read1, read2, and singleton files) were then mapped to a dark-eyed junco (*Junco hyemalis;* Passeriformes: Passerellidae) reference sequence (MLZ69236; Friis et al., 2018). Resulting sequences were indexed using BWA (Li & Durbin, 2009) and SAMtools (Li et al., 2009). Next, we used PICARD (Broad Institute, 2019) to clean the alignments, add read group header information, and remove PCR and sequencing duplicates. Single nucleotide polymorphisms (SNPs) were called for each individual against the reference, and Genome Analysis Toolkit (GATK, v.4.2.1; McKenna et al., 2010) was used to call and realign around indels, call and annotate SNPs, and filter SNPs around indels. We restricted the data to high-quality SNPs by adding a quality filter (Q30) before converting the resulting VCF file to a FASTA file using GATK.

### Selection Analyses

To determine whether genes associated with phenotype were under positive selection, we identified candidate genes and tested for signs of selection in *M. m. maxima*. Candidate genes were chosen through a review of literature that identified genes that showed correlations with phenotypes of interest. We chose 34 candidate genes associated with body size (Liu et al., 2015), bill size and shape (Lamichhaney et al., 2015; Chaves et al., 2016; Huang et al., 2022), migration (Vallone et al., 2007; Ruegg et al., 2014; Delmore et al., 2015), dispersal (San-Jose et al., 2023), plumage coloration (Walsh et al., 2011), and salt tolerance (Walsh et al., 2019). Sequences for each gene were obtained from NCBI GenBank (Table S3). Once reference sequences were obtained, we extracted corresponding sequences in our outgroup taxa *M. georgiana* and *M. m. maxima* using a custom BLAST search in Geneious Prime 2023.0.4 (www.geneious.com). Additionally, we extracted gene sequences from a *M. m. maxima* reference sequence (UAM31500; Louha et al., 2020). Sequences with no hits or had an alignment score of less than 60% were discarded. If one transcript yielded multiple hits, only the highest-scoring hit was retained for further analysis. The sequences were aligned by gene using MUSCLE (Edgar, 2004). We tested 26 candidate genes associated with body size (n = 5), bill size (n = 5), migration (n = 8), dispersal (n = 3), plumage color (n = 3), and salt tolerance (n = 2; Table 2) using the McDonald-Kreitman (MK) test, which compares the polymorphisms in one species or population to the divergence between multiple species (McDonald & Kreitman, 1991; Egea, Casillas & Barbadilla, 2008). The MK test was performed using an online tool (http://mkt.uab.es/mkt/mkt.asp; Egea, Casillas & Barbadilla, 2008).

### Phylogeny Reconstruction

To reconstruct the phylogeny of the *M. melodia* complex and outgroups, we extracted ultraconserved elements (UCEs) from the whole-genome FASTA sequence files using PHYLUCE (Faircloth, 2016). We used UCEs because they are used to capture phylogenomic information from both shallow and deep time depths (Faircloth et al., 2012). We included the *Melospiza* individuals we sampled and the *J. hyemalis* reference sequence. Individual UCE loci were aligned using MUSCLE (Edgar, 2004) and were edge trimmed. We filtered this dataset to include loci that had all the *Melospiza* individuals and the *J. hyemalis* reference represented in a 95% matrix. The sequences were aligned by loci using MUSCLE and then concatenated into one dataset using PHYLUCE (Edgar, 2004; Faircloth, 2016). We obtained branch supports with the ultrafast bootstrap approximation (Hoang et al., 2018) implemented in IQ-TREE software (v.2.1.4; Minh et al., 2020). The phylogenetic tree was visualized and edited with FigTree (v.1.4.4; http://tree.bio.ed.ac.uk/software/Figtree/)

## Results

### Phenotypic Variation

Phenotypic variation among the subspecies was pronounced. Overall, subspecies that occur in the western range (*maxima, sanaka*, and *insignis*) were larger than eastern Alaska subspecies (*rufina* and *caurina*). *Melospiza melodia maxima* was approximately 1.6 times larger in body mass than *rufina* (Table 1; Fig. S1). For all other phenotypic measurements, differences were ∼1.2 times larger (Table 1). In all measurements, there was no significant difference between *rufina* and *caurina* (Table 1). Additionally, *maxima* and *sanaka* showed differences only between bill height and bill width (Table 1).

**Table 1:** Mean (± standard deviation) for the phenotypic traits compared across the five song sparrow subspecies of interest. Values with different letters are significantly different based on a Tukey’s post-hoc test.

|  | *M. m. maxima* | *M. m. sanaka* | *M. m. insignis* | *M. m. caurina* | *M. m. rufina* |
| --- | --- | --- | --- | --- | --- |
| Mass (g) | 46.2 ± 2.71 (A) | 45.4 ± 2.06 (A) | 41.0 ± 2.61 (B) | 28.4 ± 1.77 (C) | 27.9 ± 1.75 (C) |
| Wing chord length (mm) | 82.7 ± 3.04 (A) | 83.3 ± 2.45 (A) | 80.9 ± 2.14 (A) | 72.6 ± 1.43 (B) | 69.7 ± 3.03 (B) |
| Tail length (mm) | 78.9 ± 4.27 (A) | 77.8 ± 4.28 (A) | 75.4 ± 4.52 (A) | 67.1 ± 4.28 (B) | 65.8 ± 3.01 (B) |
| Tarsus length (mm) | 27.6 ± 2.04 (A) | 27.0 ± 0.99 (A) | 26.3 ± 0.53 (AB) | 24.2 ± 1.32 (BC) | 24.2 ± 1.41 (C) |
| Bill length (mm) | 12.4 ± 1.34 (A) | 12.4 ± 0.51 (A) | 11.9 ± 0.57 (AB) | 10.7 ± 0.60 (B) | 10.8 ± 2.75 (B) |
| Bill height (mm) | 7.5 ± 1.08 (A) | 6.89 ± 1.04 (B) | 6.5 ± 0.22 (AB) | 6.1 ± 0.45 (AB) | 6.37 ± 0.87 (B) |
| Bill width (mm) | 5.9 ± 0.77 (A) | 5.6 ± 0.56 (B) | 5.3 ± 0.32 (BC) | 5.1 ± 0.34 (BC) | 5.1 ± 0.45 (C) |
| Skull length (mm) | 38.8 ± 1.54 (A) | 38.8 ± 0.80 (A) | 38.0 ± 0.95 (A) | 35.7 ± 1.93 (B) | 34.8 ± 1.41 (B) |

PCA analysis revealed that 55% of the variation among the five subspecies of *M. melodia* was explained in PC1 by five phenotypic traits: mass, wing chord, tail length, tarsus length, and skull length. Around 13% of the variation was explained in PC2 by the remaining three traits: bill length, bill width, and bill height (Fig. 3). In the PCA graph, individuals tend to clump into two groups, mostly divided along PC1. The three western subspecies (*maxima, sanaka,* and *insignis*) group together, and the two remaining eastern subspecies (*caurina* and *rufina*) also group together (Fig. 3).

**Figure 3:**
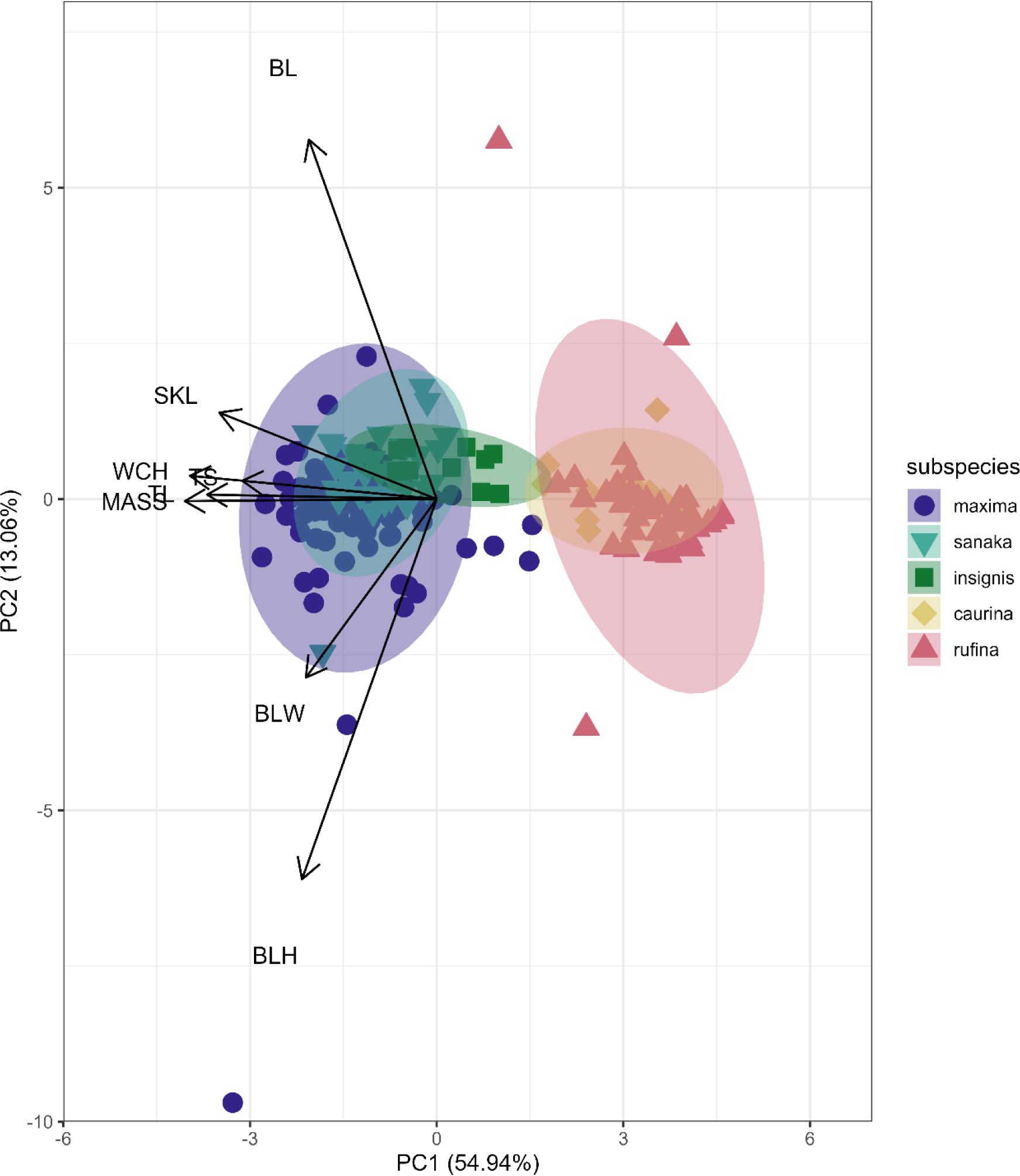
PCA plot of phenotypic data for the five *Melospiza melodia* subspecies. PC1 explains approximately 55% of the variation with five phenotypic traits: mass (MASS), wing chord (WCH), tail length (TL), tarsus length (TS), and skull length (SKL) length. PC2 explains 13% of the variation with the three remaining traits: bill length (BL), bill width (BLW), and bill height (BLH). Three subspecies (*maxima, sanaka*, and *insignis*) group together and are larger from the second group of two subspecies (*caurina* and *rufina*). Colors correspond to Figure 1.

### Summary Statistics of Whole-genome Sequencing

We obtained >400 million reads, ranging from 12,566,699 to 84,504,717 per individual (average = 60,119,612; Table S4). Coverage averaged 6.7x, ranging from 2.1x to 9.7x per individual (Table S4).

### Selection Analyses

Of the 26 candidate genes tested for signals of positive selection, two showed significance: *BCO1* (p = 0.037) and *KCTD21* (p = 0.032; Table 2). These genes are associated with plumage color and dispersal, respectively (Ruegg et al., 2014; Toews, Hofmeister & Taylor, 2017; San-Jose et al., 2023).

**Table 2:** *p*-value results from McDonald-Kreitman tests. Gene names and p-values in bold denote significant results (< 0.05). Significance indicates a gene is under positive selection.

| Phenotypic Trait | Candidate Genes | MK Test *p*-values | Citation |
| --- | --- | --- | --- |
| Bill size | *ALX1* | NULL | Lamichhaney et al., 2015 |
|  | *CCDC149* | 0.781 | Huang et al., 2022 |
|  | *DLK1* | 0.611 | Chaves et al., 2016 |
|  | *HMGA2* | 0.145 | Chaves et al., 2016 |
|  | *LGI2* | 0.280 | Huang et al., 2022 |
| Body size | *AFAP1* | 0.376 | Liu et al., 2015 |
|  | *GGA1* | 0.808 | Liu et al., 2015 |
|  | *LCORL* | NULL | Liu et al., 2015 |
|  | *QDPR* | 0.363 | Liu et al., 2015 |
|  | *SLIT2* | 0.792 | Liu et al., 2015 |
| Plumage color | *APOD1* | 0.316 | Walsh et al., 2011 |
|  | ***BCO1*** | **0.037** | Walsh et al., 2011 |
|  | *CD36* | 0.924 | Walsh et al., 2011 |
| Dispersal | ***KCTD21*** | **0.032** | San-Jose et al., 2023 |
|  | *SLC2A1* | NULL | San-Jose et al., 2023 |
|  | *TGFB2* | 0.264 | San-Jose et al., 2023 |
| Migration | *CLOCK* | 0.849 | Vallone et al., 2007 |
|  | *CREB1* | 0.820 | Ruegg et al., 2014 |
|  | *CRY1* | NULL | Vallone et al., 2007 |
|  | *CRY2* | 0.675 | Vallone et al., 2007 |
|  | *NPAS3* | 0.300 | Ruegg et al., 2014 |
|  | *PER2* | 1.000 | Vallone et al., 2007 |
|  | *PER3* | 0.314 | Ruegg et al., 2014 |
|  | *YPEL1* | 0.279 | Delmore et al., 2015 |
| Salt tolerance | *MMP17* | 0.156 | Walsh et al., 2019 |
|  | *SLC9A3* | 0.868 | Walsh et al., 2019 |

### Phylogenetic Reconstruction

After filtering UCE loci present in the *Melospiza* and *J. hyemalis* genomes to obtain a 95% matrix, the final alignment contained 4793 loci. All nodes within the *M. melodia* clade received 100% support from the ultrafast bootstrap (Fig. 4). The phylogenetic tree shows that two of the western subspecies (*sanaka* and *insignis*) and the two eastern subspecies (*caurina* and *rufina*) form their own clades. However, *maxima* is a sister group to all other *M. melodia* subspecies (Fig. 4).

**Figure 4:**
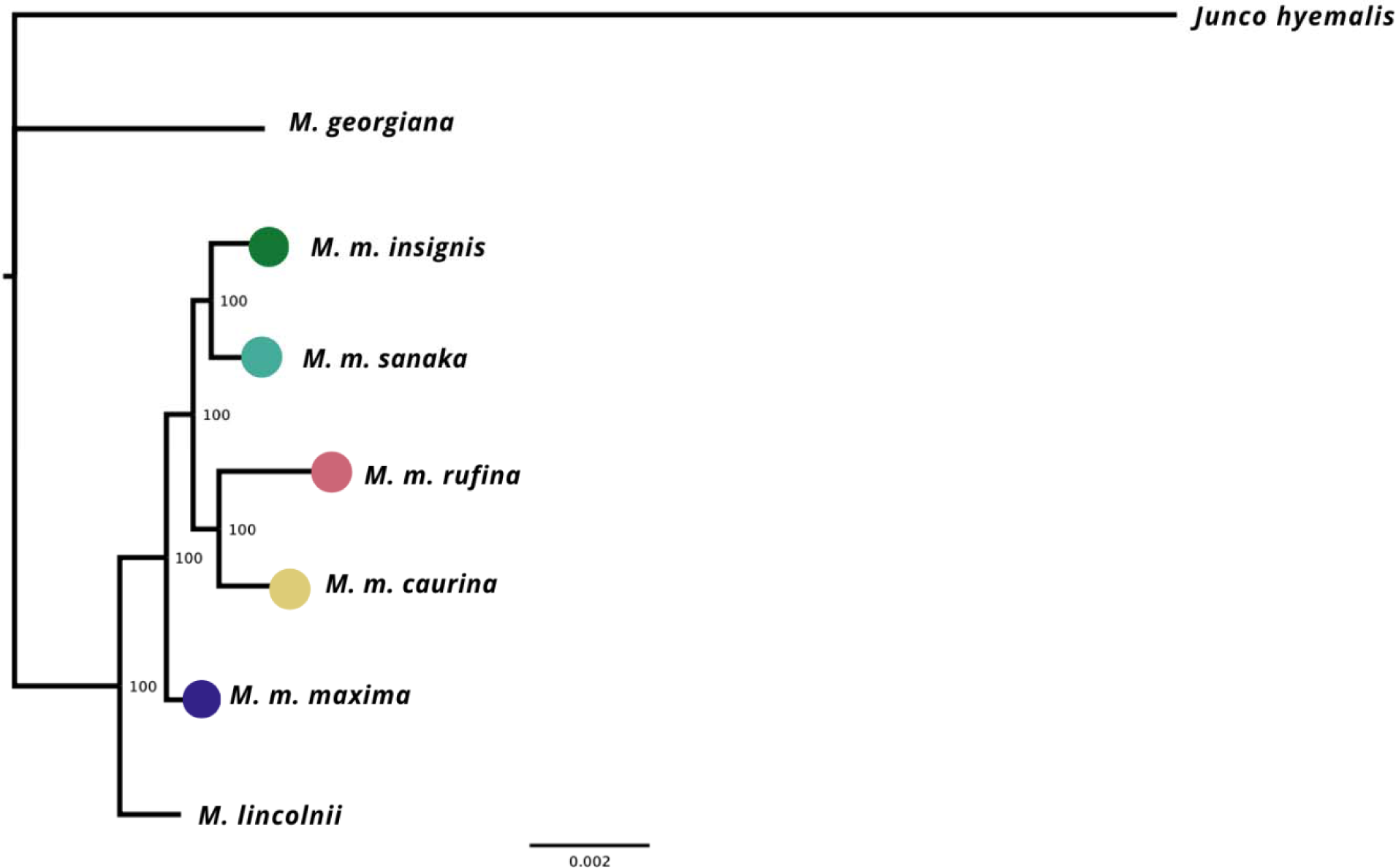
Maximum-likelihood phylogeny based on 4793 UCE loci. Values on the nodes represent bootstrap values from 1000 iterations. *M. m. maxima* is shown to be a sister group to the other *M. melodia* subspecies.

## Discussion

Our results show that the westernmost subspecies of song sparrow (*M. m. maxima*) is approximately 1.6 times larger than eastern *rufina* (Table 1). We found evidence for positive selection in two genes (*BCO1* and *KCTD21*) in *M. m. maxima* (Table 2). We reconstructed a novel phylogeny for the Alaska song sparrow complex, which places *M. m. maxima* outside the rest of the *M. melodia* clade (Fig. 4). This suggests that song sparrows colonized the Aleutian Islands early and underwent divergence before prior to the eastern portion of the species’ current Alaska range was recolonized.

### Phenotypic Variation

The strong increase in body size among resident populations in the western Aleutians (Table 1) is concordant with previous research (Aldrich, 1984) and follows Bergmann’s Rule, by which endothermic animals tend to be larger in cooler climates (Bergmann, 1847, Meiri & Dayan, 2003; Carbeck et al. 2023). When examining the PCA results (Fig. 3), the three western subspecies (*maxima, sanaka*, and *insignis*) grouped together and the remaining two eastern subspecies (*caurina* and *rufina*) grouped together. These groups are consistent with the different life-history strategies, with the larger, western subspecies being sedentary and the smaller, eastern subspecies being migratory. Sedentary birds tend to comply with Bergmann’s rule more than migratory species because migratory species can seasonally escape more severe climatic conditions (Meiri & Dayan, 2003; but see Ashton, 2002). Likewise, island living could also have an impact on body size. Birds and mammals experience the “island rule”, whereby large organisms on islands evolve toward smaller sizes and small organisms evolve toward larger sizes (Losos & Ricklefs, 2009).

*Melospiza melodia maxima* and *sanaka* were similar in all measurements except for bill height and bill width (Table 1). *M. m. sanaka* had significantly narrower and shallower bills than *maxima,* which is consistent with previous research (Patten & Pruett, 2009; Gibson & Withrow, 2015). Slender bills are associated with invertebrate consumption, while stouter bills are useful for eating seeds (Aldrich, 1984). However, current knowledge does not include differences in diet between these two subspecies that might explain this difference in morphology (Aldrich, 1984). It is possible that the thicker bills of *maxima* were an adaptation for living in a glacial refugium during the Last Glacial Maximum, but the exact reason is unknown.

### Selection Analyses

We found two genes of 26 that showed signs of positive selection: *BCO1* and *KCTD21* (Table 2). *BCO1* is associated with coloration and carotenoid breakdown (Toews, Hofmeister & Taylor, 2017). Carotenoids, which are obtained from an animal’s diet, must be metabolized to be incorporated into tissues. Beta-carotene (β-carotene) oxygenases (i.e., *BCO1* and *BCO2*) are some of the best-known genes involved in carotenoid breakdown. *BCO1* facilitates the separation of vitamin A from β-carotene, whereas *BCO2* is more directly involved in coloration (Eriksson et al., 2008; Toews, Hofmeister & Taylor, 2017). After carotenoid breakdown, the pigments can be deposited into feathers and skin. Many studies examine the expression of these candidate genes in colorful birds. For example, Walsh et al. (2011) found that *BCO1* was expressed in developing feathers and skin of queleas, another bird species. However, there are many different strategies for carotenoid deposition and numerous genes associated with carotenoid coloration, some that we did not include in our study design (Eriksson et al., 2008; Walsh et al. 2011).

Potassium channel tetramerization domain, *KCTD21*, is associated with dispersal, migration, and the circadian clock (San-Jose et al., 2023). Dispersal and migration are two separate, but possibly intertwined, processes. Migration is the seasonal go-and-return movements between breeding and non-breeding grounds, whereas dispersal is the movement from the place of birth to breed somewhere else (Dingle & Drake, 2007). In a study of the genetic underpinnings of dispersal in reptiles, San-Jose et al. (2023) found that multiple genes associated with the circadian clock and migration were differently expressed in dispersers. This suggests that some of the same processes are involved in migration and dispersal. We suspect this gene remains under positive selection due to ancestral *maxima* individuals dispersing to the western Aleutian Islands.

We expected to see more candidate genes under positive selection in *maxima*, such as genes related to body size and salt tolerance. For example, a recent study by Carbeck et al. (2023) found evidence of natural selection acting on genes associated with body size. The authors found nine candidate genes associated with body size showed divergence between *maxima* and *rufina* song sparrows and then validated these findings by predicting the genotypes of five small song sparrows in California (Carbeck et al., 2023). Our study did not test for selection any candidate genes that Carbeck et al. (2023) found. Further work with these candidate genes could determine if these genes are under adaptive selection in *maxima*.

### Phylogenetic Reconstruction

Our phylogenetic tree shows that the westernmost subspecies *maxima* is a sister taxon to all other Alaska *M. melodia* subspecies (Fig. 4). Further, the eastern Alaska subspecies *caurina* and *rufina* are grouped together and two of the western subspecies (*sanaka* and *insignis*) form another clade (Fig. 4). This suggests that *maxima* colonized the western Aleutian Islands first, underwent differentiation from other forms, and then at some later time, the eastern part of the Alaskan range was colonized by ancestors of the remaining subspecies. One possibility for this colonization history is that following an initial colonization of Alaska, *maxima* survived in an Aleutian glacial refugium (Pruett & Winker, 2005a; Winker et al., 2023) while other Alaska populations were extirpated, and then Alaska was re-colonized post-glacially. This hypothesis is inconsistent with the colonization history postulated by Pruett and Winker (2005b), which suggested a probable postglacial stepwise process from east to west.

Interestingly, *maxima* and *sanaka* are similar in size and have similar life-history patterns, even though *M. m. sanaka* is more closely related to all of the non-*maxima* subspecies. This phenotypic similarity could be due to convergent evolution in similar environments, or, more likely, due to adaptive alleles for Aleutian life introgressing through gene flow from *maxima* through (Pruett & Winker, 2005b; Reding et al., 2008; Graham et al., 2021).

It is possible that gene flow could be obscuring our phylogenetic results, however we do not expect this to be the case. Models show that moderate to high levels of gene flow can degrade phylogenetic signal and thus produce inaccurate phylogenetic trees (Eckert & Carstens, 2008; Leaché et al., 2014). Previous research by Pruett and Winker (2005b) found gene flow between *maxima* and *sanaka* to be at approximately 1.20 – 2.90 individuals per generation (Fig. S2). The exact amount of gene flow needed to obscure phylogeny is not noted but the current estimates for gene flow in the Aleutian Islands are not very high. Future research should determine if this obscuration is happening within this song sparrow complex.

## Conclusion

Among song sparrows in Alaska, phenotypic differences between subspecies suggest that selective pressures have acted heterogeneously on different populations. We found two candidate genes under positive selection in *M. m. maxima*. Genes under selection associated with carotenoid breakdown and dispersal (*BCO1* and *KCTD21*, respectively), suggest that *maxima* has undergone selective pressure for these traits relative to other members of the species in this region. The phylogenetic relationship that we recovered among Alaska song sparrow subspecies suggests a new hypothesis for the species’ biogeographic history. We suggest that ancestral populations of *M. m. maxima* colonized Alaska and the Aleutian Islands, and then persisted in a glacial refugium, likely while other populations were extirpated. Then after a substantial period of time the rest of the state was recolonized by ancestors of the remaining subspecies. This complex might be an ideal candidate for future studies of evolution of island populations and local adaptation.

## Acknowledgements

We thank Jacob Adams, Robin Andrews, Syndonia Bret-Harte, Stefanie Ickert-Bond, Rachel Richardson, Alex Sletten, Laura Weingartner, and Jeff Wells for their invaluable input on earlier drafts. We also want to thank all museum collectors who collected the specimens used in this project.

## Supplement

**Supplemental Table S1: Accession numbers and genus, species, and subspecies used in phenotypic analysis.** UAM = University of Alaska Museum

**Supplemental Table S2: Sample information used in the genomic analysis.** UAM = University of Alaska Museum. All sequences generated in this study are archived under SRA project *pending*. † denotes individual used as a reference sequence.

**Supplemental Table S3: Accession numbers, gene names, and associated phenotypic traits for sequences that were tested in gene selection.** Asterisk (*) denotes sequences dropped from analysis due to poor BLAST results.

**Supplemental Table S4: Summary statistics of sequenced datasets including total number of reads, percentage of properly paired reads, and mean depth of coverage.**

**Supplemental Figure S1: Boxplots of phenotypic data for each *Melospiza melodia* subspecies:** a) mass (g), b) wing chord (mm), c) tail length (mm), d) tarsus length (mm), e) bill length (mm), f) bill height (mm), g) bill width (mm), and h) skull length (mm). Colors correspond to the colors in Figure 1.

**Supplemental Figure S2: Map of song sparrow range in Alaska with pairwise estimates of directional gene flow (genetic migration, *N_e_m*) from Pruett & Winker (2005b).** Values denote the number of migrant individuals per generation relative to effective population size, and the arrows denote direction of migration. For example, *M. m. maxima* is receiving 2.90 migrants per generation from *M. m. sanaka*. Colors correspond to Figure 1.

