## Supplementary Materials for "Evidence of positive selection and a novel phylogeny among five subspecies of song sparrow (*Melospiza melodia*) in Alaska"

Supplemental Table S2: Sample information used in the genomic analysis. UAM = University of Alaska Museum. All sequences generated in this study are archived under SRA BioProject PRJNA1114297. † denotes individual used as a reference sequence.

| Species | Locality | Catalogue Number | Tissue Number | Sequence Archive |
| --- | --- | --- | --- | --- |
| *Junco hyemalis*† | Virginia: Giles County, Mount Lake Biological Station | MLZ:Bird:69236 | n.a. | PRJNA493001 |
| *Melospiza georgiana* | Wisconsin: Burnett County, Saint Croix National Riverway | UAM11868 | CLP550 | SAMN41480621 |
| *Melospiza lincolnii* | Alaska: Southeast, Gravina Island | UAM21920 | ABJ048 | SAMN41480622 |
| *Melodia melodia maxima* | Alaska: Aleutian Islands, Attu Island | UAM31500 | UAMX4047 | PRJNA511035 |
| *Melospiza melodia maxima* | Alaska: Aleutian Islands, Adak Island | UAM10946 | CLP108 | SAMN41480625 |
| *Melospiza melodia sanaka* | Alaska: Shumagin Islands, Popof Island | UAM11585 | CLP251 | SAMN41480627 |
| *Melospiza melodia insignis* | Alaska: Kodiak Island, Cape Chiniak | UAM14002 | DDG1900 | SAMN41480624 |
| *Melospiza melodia caurina* | Alaska: Copper River Delta | UAM11384 | CLP020 | SAMN41480623 |
| *Melospiza melodia rufina* | Alaska: Southeast, Hyder | UAM7343 | KSW1374 | SAMN41480626 |

Supplemental Table S3: Accession numbers, gene names, and associated phenotypic traits for sequences that were tested in gene selection. Asterisk (*) denotes sequences dropped from analysis due to poor BLAST results.

| Associated Phenotypic Trait | Accession Number | Gene |
| --- | --- | --- |
| Bill size | XM_002196007.3 | *ALX1* |
| Bill size | XM_030270858.3 | *CCDC149* |
| Bill size | XM_030274996.3 | *DLK1* |
| Bill size | XM_004176091.5 | *HMGA2* |
| Bill size | XM_041715714.1 | *LGI2* |
| Body size | XM_030270679.3 | *AFAP1* |
| Body Size | XM_002198985.6 | *GGA1* |
| Body size | XM_030270815.3 | *LCORL* |
| Body size | XM_002193889.6 | *QDPR* |
| Body size | XM_030270827.3 | *SLIT2* |
| Body size | XM_030276710.3 | *WAPL** |
| Body size | XM_002193767.6 | *LAP3** |
| Plumage color | XM_015637109.2 | *APOD1* |
| Plumage color | XM_032750540.2 | *BCO1* |
| Plumage color | XM_030291411.3 | *CD36* |
| Plumage color | XR_003962894.3 | *SCARB1** |
| Plumage color | XM_030273867.3 | *HPS5** |
| Plumage color | XM_030256157.3 | *STARD3** |
| Plumage color | XM_030290911.3 | *BCO2** |
| Dispersal | XM_012569888.4 | *KCTD21* |
| Dispersal | XM_004175099.5 | *TGFB2* |
| Dispersal | XM_032752076.2 | *SLC2A1* |
| Migration | XM_030271098.3 | *CLOCK* |
| Migration | XM_030276951.3 | *CREB1* |
| Migration | XM_030263133.3 | *CRY1* |
| Migration | XM_012574176.4 | *CRY2* |
| Migration | XM_032748493.2 | *NPAS3* |
| Migration | XM_030280764.3 | *PER2* |
| Migration | XM_030289516.3 | *PER3* |
| Migration | XM_030285787.3 | *YPEL1* |
| Salt tolerance | XM_041719360.1 | *MMP17* |
| Salt tolerance | XM_030265641.3 | *SLC9A3* |
| Salt tolerance | XM_030276092.3 | *MYOF** |
| Salt tolerance | XM_030283214.3 | *WNK2** |

Supplemental Table S4: Summary statistics of sequenced datasets including total number of reads, percentage of properly paired reads, and mean depth of coverage.

| Taxa | Tissue Number | Number of reads | % properly paired reads | Mean depth of coverage |
| --- | --- | --- | --- | --- |
| *Melospiza georgiana* | CLP550 | 81,339,509 | 93.0% | 7.04 |
| *Melospiza lincolnii* | ABJ048 | 12,566,699 | 92.9% | 2.10 |
| *Melospiza melodia maxima* | CLP108 | 26,989,233 | 91.4% | 3.33 |
| *Melospiza melodia sanaka* | CLP251 | 84,504,717 | 94.0% | 9.71 |
| *Melospiza melodia insignis* | DDG1900 | 73,078,606 | 92.0% | 8.62 |
| *Melospiza melodia caurina* | CLP020 | 69,765,460 | 92.0% | 8.12 |
| *Melospiza melodia rufina* | KSW1374 | 72,593,062 | 90.6% | 8.08 |

Supplemental Figure S1: Boxplots of phenotypic data for each *Melospiza melodia* subspecies: a) mass (g), b) wing chord (mm), c) tail length (mm), d) tarsus length (mm), e) bill length (mm), f) bill height (mm), g) bill width (mm), and h) skull length (mm). Colors correspond to the colors in Figure 1.

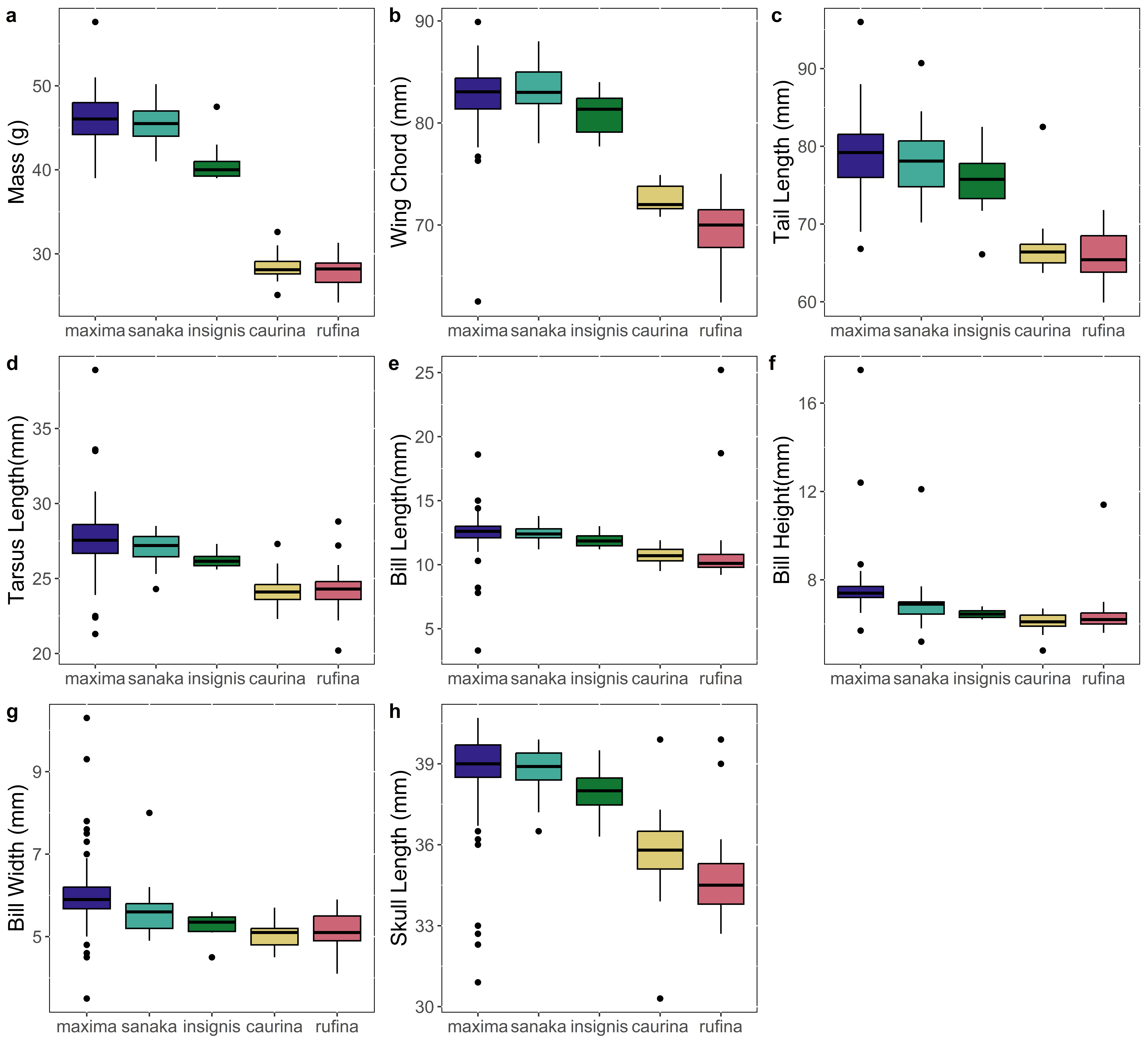

Supplemental Figure S2: Map of Song Sparrow range in Alaska with pairwise estimates of directional gene flow (genetic migration, *N_e_m*) from Pruett & Winker (2005b). Values denote the number of migrant individuals per generation relative to effective population size, and the arrows denote direction of migration. For example, *M. m. maxima* is receiving 2.90 migrants per generation from *M. m. sanaka*. Colors correspond to Figure 1.

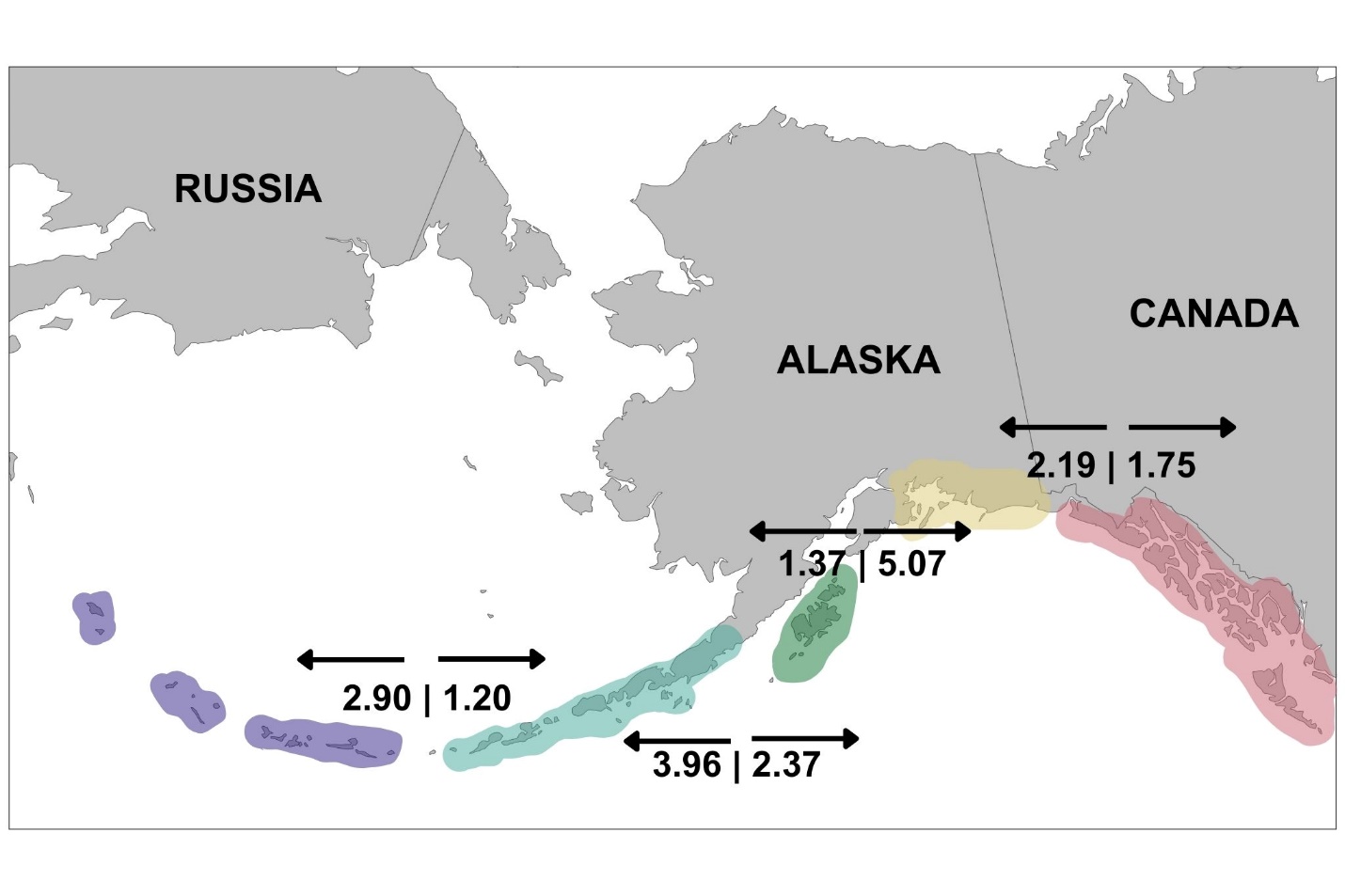
